## Supplementary figures for "*Sox9* prevents retinal degeneration and is required for limbal stem cell differentiation in the adult mouse eye": SupplementaryFigures.pdf

**By**

Alicia Hurtado, Víctor López-Soriano, Miguel Lao, M. Ángeles Celis-Barroso, Pilar Lazúen, Alejandro Chacón de Castro, Yolanda Ramírez-Casas, Miguel Alaminos, J. Martin Collinson, Miguel Burgos, Rafael Jiménez, F. David Carmona, Francisco J Barrionuevo

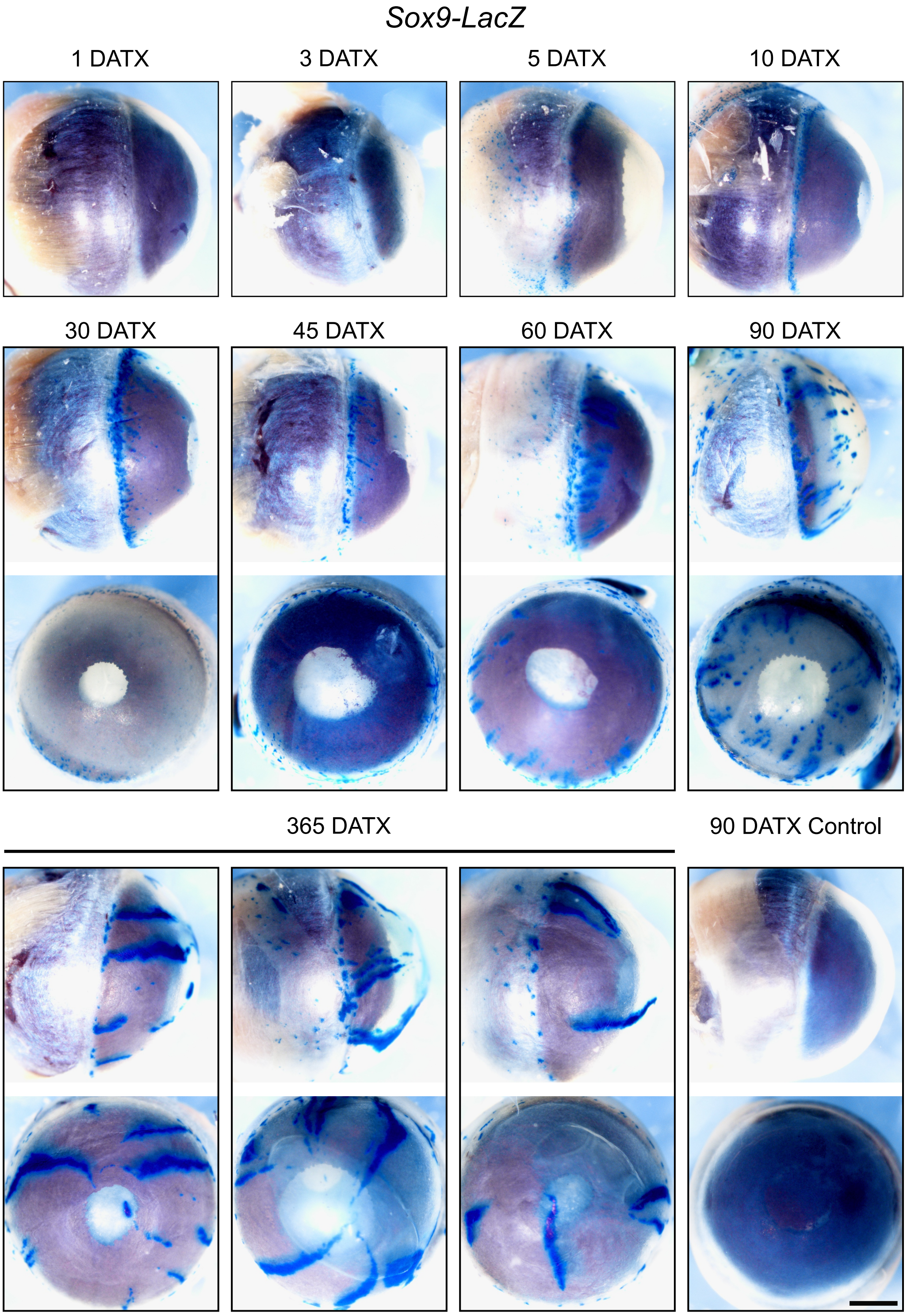


**Figure S1**. Whole mount X-gal staining of control and *Sox9-Lac*Z eyes at different days after tamoxifen (DATX) administration. Scale bar represents 1 mm.

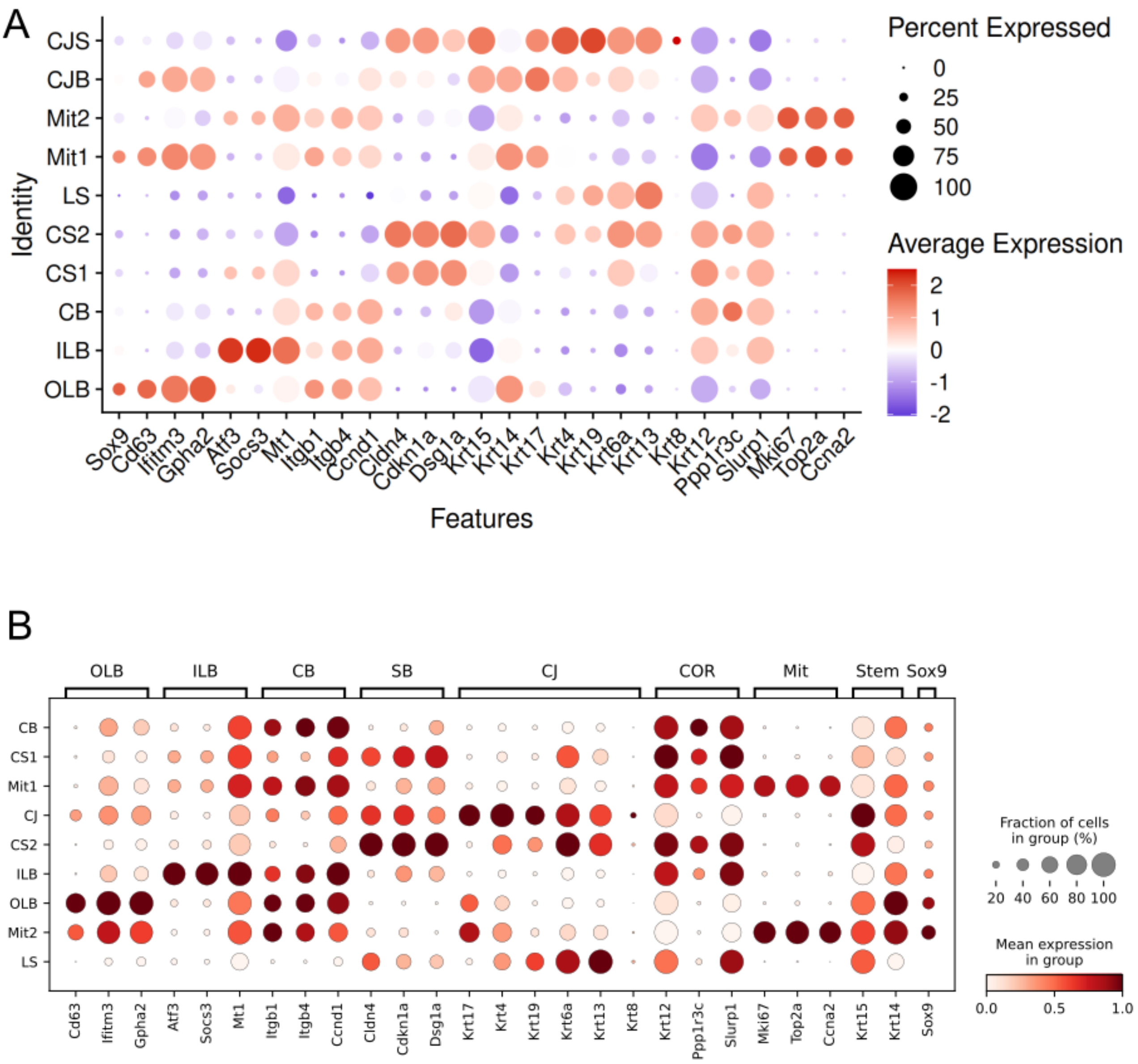


**Figure S2.** Dot plot showing the expression of specific markers in the cell groups identified from the single-cell RNA-seq dataset of isolated limbal epithelial cells reported by Altshuler et al. (2022). **(A)** Analysis using the Seurat R package. **(B)** Analysis using the Scanpy Python package.

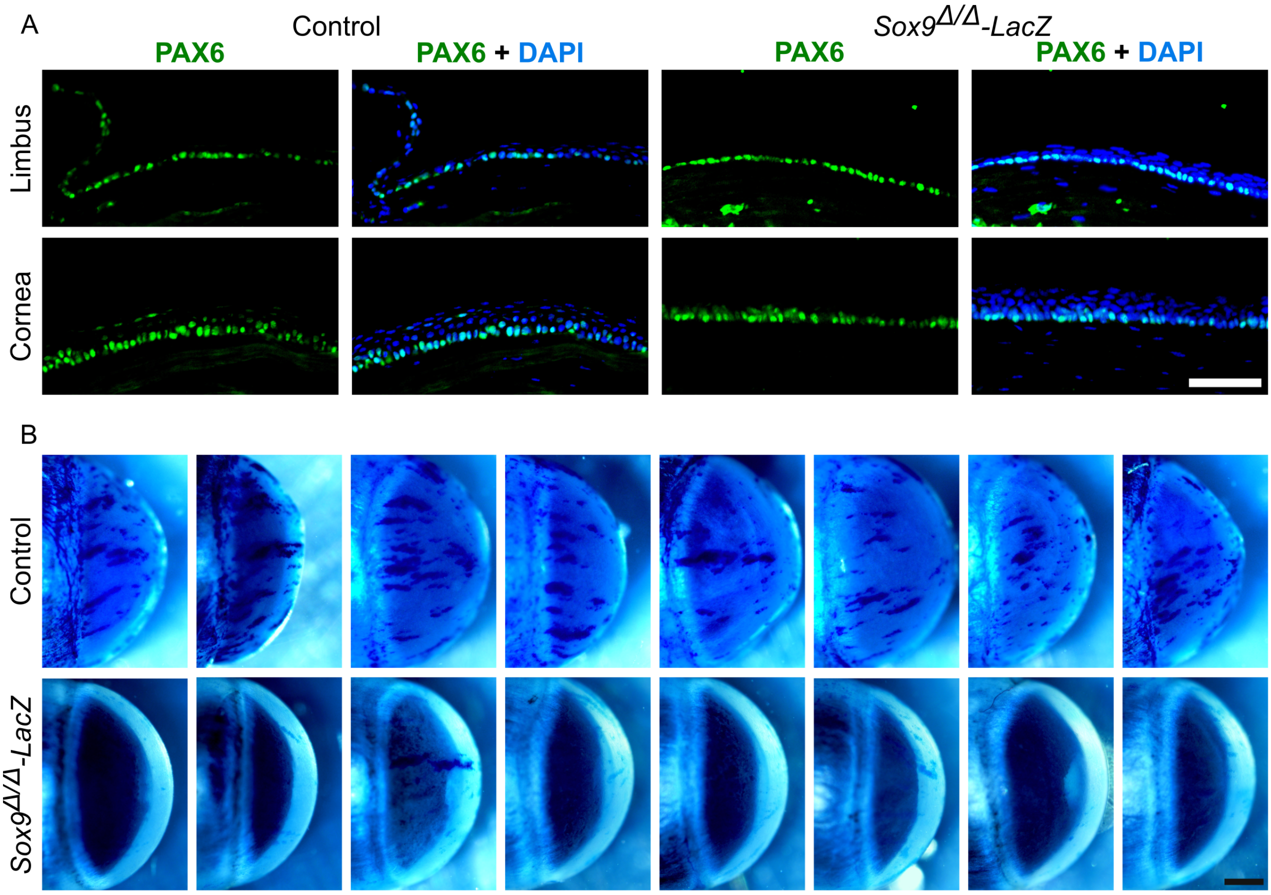


**Figure S3.** Effect of *Sox9* deletion on PAX6 expression and stem cell differentiation in the limbus and cornea. **(A)** Immunofluorescence staining for PAX6 in the limbus and cornea from control and *Sox9*$^{\Delta/\Delta}$ mice. DAPI (blue) was used to counterstain nuclei. **(B)** X-gal staining of all analyzed eyes from control and *Sox9*$^{\Delta/\Delta}$-LacZ mice at 98 days after tamoxifen administration. Scale bars represent 500 µm.
